# Predicting microbial community compositions in wastewater treatment plants using artificial neural networks

**DOI:** 10.1101/2022.09.08.507071

**Authors:** Xiaonan Liu, Yong Nie, Xiao-Lei Wu

## Abstract

Activated sludge (AS) of wastewater treatment plants (WWTP) is one of the world’s largest artificial microbial ecosystems and the microbial community of the AS system is closely related to WWTP performance. However, how to predict its community structure is still unclear. Here, we used artificial neural networks (ANN) to predict the microbial compositions of AS systems collected from WWTPs located worldwide. We demonstrated that the microbial compositions of AS systems are predictable using our approach. The predictive accuracy R^2^_1:1_ of Shannon-Wiener index reached 60.42%, and the average R^2^_1:1_ of ASVs appearing in at least 10% of samples (ASVs_>10%_) and core taxa were 35.09% and 42.99%, respectively. We also found that the predictability of ASVs_>10%_ was significantly positively correlated with their relative abundance and occurrence frequency, but significantly negatively correlated with potential migration rate. The typical functional groups such as nitrifiers, denitrifiers, polyphosphate-accumulating organisms (PAOs) and glycogen-accumulating organisms (GAOs), and filamentous organisms in AS systems could also be well recovered using an ANN model, with the R^2^_1:1_ ranging from 32.62% to 56.81%. Furthermore, we found that industry wastewater source (IndConInf) had good predictive abilities, although its correlation with ASVs_>10%_ in the Mantel test analysis was weak, which suggested important factors that cannot be identified using traditional methods may be highlight by the ANN model. Our results provide a better understanding of the factors affecting AS communities through the prediction of the microbial community of AS systems, which could lead to insights for improved operating parameters and control of community structure.

## Introduction

With the increasing expansion of urbanization, about 360 billion m^3^ of wastewater is produced every year globally (1). The activated sludge (AS) system in wastewater treatment plants (WWTP) is at the heart of current sewage treatment technology(2). Microorganisms treat almost 60% of this wastewater in AS systems before release(3). This process relies on the degradation of organic compounds, biotransformation of toxic substances, and removal of pathogens by diverse microorganisms (4-6). Thus, the microbial communities present in these systems determine their performance (7). Predicting the microbial communities of AS systems and exploring factors that influence them will provide reasonable suggestions for the design, optimization, and stable operation of sewage treatment systems (8-11). However, because wastewater always contains a multiplicity of resources, the AS system exhibits an enormous microbial diversity and varies greatly worldwide. The global activated sludge community encompass about 1 billion bacterial phylotypes, with a small global core bacterial community consisting of only 28 OTUs(3). It is still unclear how we can predict such overwhelming diverse microbial communities in the AS systems of WWTPs according to the design parameters and environmental data.

AS system contains high biomass and microbial diversity(3, 12), and predicting the microbial community is complicated by diverse factors. For example, the type of wastewater directly impacts microbial composition. For example, AS systems treating municipal and industrial wastewater harbor distinct microbial communities (13, 14). The influent biodegradability [biological oxygen demand/chemical oxygen demand (B/C ratio)] also plays an essential role in shaping the AS microbial community. Low or high B/C ratio may lead to low microbial diversity and pollutant removal loading(15). Recently, the integration of high-throughput sequencing and multivariate statistical analysis indicated that the microbial communities of AS systems are correlated with multiple factors, such as location, geographical distance, dissolved oxygen (DO), temperature, hydraulic retention time (HRT), sludge retention time (SRT), inflow and effluent of chemical oxygen demand (COD), total nitrogen (TN), total phosphorus (TP) (16, 17). The combined influence of multiple environmental factors has made it difficult to predict the microbial compositions in AS systems using current methods, and thus has become an obstacle to guiding the operation of WWTPs.

Multiple linear regression models can predict microbial community structure from multiple environmental factors. A previous study predicted bacterial and fungal groups in a soil microbial community from typical soil environmental factors [C and N concentrations, pH, mean annual temperature (MAT), mean annual precipitation (MAP) and net primary productivity (NPP), etc.] using this method(18). However, since multiple regression models ignore the interaction effects of environmental factors and nonlinear relationships, the predictability of the microbial taxa in that study was at most no more than 60%. The AS system is affected by multiple cross-complex factors, including geographical factors, design and operation parameters and physicochemical parameters, and multiple regression analysis is not enough to capture this complex relationship.

Artificial neural network (ANN) is a machine learning method for the automatic and quantitative learning of a suitable relationship and providing guidance for system optimization, which is an ideal alternative to model these complex relationships between microbial communities and environmental variables as this method is better suited to account for the non-linear associations between variables and the interactions among predictors(19). ANNs have helped researchers to successfully analyze the relationship between environmental factors and microbial community structure in many ecosystems(20-22), while the relevant applications of activated sludge systems are still lacking. In addition, in previous studies, no attention has been paid to the regularity of predictability of microbial taxa, which is essential for a deeper understanding and control of microbial community structure.

Here, we used ANN models and environment data to predict the microbial community structure of AS systems from global wastewater treatment plants. We analyzed the predictability of different taxa and the effects of environmental factors on the prediction. These analyses deepened our understanding of the microbial community of AS systems, provided reasonable suggestions for accurately predicting major functional groups, and provided a theoretical basis for better design and operating parameters, and to control community structure.

## Methods

### Datasets

This study used a previously published dataset of 1186 activated sludge samples taken from 269 WWTPs in 23 countries across 6 continents. In addition to 16S rRNA sequencing data of these sludge samples, associated metadata conforming to the Genomic Standards Consortium’s MIxS and Environmental Ontology Standards(23) were also provided by plant managers and investigators.

Among the 1186 samples collected in the previous study, some were from different sampling points of the same WWTP, and some were obtained from the same sampling point at different times. As such, the environmental factors and community structure between these samples may have high similarities(3) and when evaluating a model with all 1186 samples, it may overestimate the generalization ability of its predictions. Therefore, we removed the samples with the same environmental information and minimal weighted-UniFrac distance (no more than Q1-3*IQR, Q1 is the first quartile, and IQR is Inter-Quartile Range) of the microbial community in these 1186 original samples and used the remaining 777 samples (no data leakage) for subsequent construction and evaluation of the predictive model.

### Data preprocessing

To ensure the accuracy, completeness, and consistency of the data, we preprocessed the original data before building the machine learning predictive model.

### Metadata preprocessing

For the metadata obtained from previous studies (reference 3), to reduce the redundancy of environmental data, we first removed some non-numerical variables of multiple categories that are difficult to operate and some variables with no practical meanings, such as site name, city name, etc. The remaining variables were used to train the model and their abbreviations and meanings are shown in Table S1. To have a clearer understanding of the environmental factors, we classified the different types of environmental factors used for prediction(3), including climate conditions, design and operation parameters, inflow conditions, effluent conditions and physicochemical properties of samples (Table S1). Then, the *LabelEncoder* algorithm was used to numericize binary non-numeric variables, and missing values were completed according to the two-nearest neighbor principle. Additionally, all environmental factors were normalized to 1-100 to eliminate dimensional influence(20).

The final environment data for input in our machine learning predictive models is shown in Table S2.

### Sequencing data preprocessing

The original microbial sequencing data were processed using Quantitative Insights Into Microbial Ecology (QIIME2) software (http://qiime2.org)(24). All paired reads were merged, quality filtered, then denoised through DADA2 plugin(25) to clustered into 100% amplicon sequence variants (ASVs). Then, ASVs classified as fungi, ASVs with unassigned taxonomy at the domain level, and ASVs annotated as mitochondria or chloroplasts were removed, so that only bacterial and archaeal sequences were retained. Singletons (ASVs with only one sequence) were discarded before further analysis to reduce the impact of sequencing errors. Then we rarefied each sample to 20954 sequences, to obtain the maximal observation of both samples and features, from which 46628 ASVs were obtained. The final feature sequences were taxonomically classified using the MiDAS4 reference database(26), and phylogenetic trees were generated using phylogeny plugins for further analysis.

Alpha diversity indices such as the Shannon-Wiener index, ASV count (species richness), and Pielou’s evenness were calculated using the vegan package of R 4.0.3 software (http://www.r-project.org) according to the final feature table. Faith’s phylogenetic diversity was calculated using the Picante package of R 4.0.3 software according to the feature table and phylogenetic tree. The relative abundance of each ASV was also calculated from this feature table. Together, these microbial community features served as target variables for our WWTP community predictive models.

### Artificial neural network model

We employed the PyTorch (v1.7.1, https://pytorch.org/) library in python 3.8 to build fully connected artificial neural networks (ANN). After testing, the three-layer network (including a hidden layer), with *relu* and *sigmoid* as activation functions between layers, had an excellent prediction effect on the condition that the network topology was relatively simple.

The first layer was the input layer, and this study’s input data was the sewage treatment plants’ environmental data (Table S2). Therefore, there were 48 nodes in the first layer. According to previously studied algorithm optimization results (27), the internal hidden layer had 97 nodes (2n+1, where n is the number of input nodes). Meanwhile, the output layer had 1 node, corresponding to the index of alpha-diversities, the relative abundance of different ASVs, or the abundance of functional groups (Fig. 2a). We built predictive models separately for each target to minimize prediction errors.

There were many random operations in the model training process, which made the results inconsistent after repeatedly running the same code. We set a fixed global seed for the random number generator to obtain repeatable training results. All models were trained twenty times by different seeds to avoid the deviation caused by each randomization, and the averaged results were used for further analysis.

### Alpha diversity and microbial taxa abundance predictive model

For alpha diversity, we established predictive models for the Shannon-Wiener index, Pielou’s evenness index, species richness, and Faith’s phylogenetic diversity. For taxa, the relative abundance of taxa with low occurrence frequency was zero in many samples, which made it difficult for the model to learn useful information on the training set (underfitting). Therefore, only the relative abundance of ASVs present in at least 10% of samples (named ASVs_>10%_, corresponding ASVs_<10%_ represent ASVs that appears in no more than 10% samples) were modeled to predict.

There were 777 samples to build the alpha diversity or ASVs_>10%_ abundance predictive models. To reduce the risk of overfitting during hyperparameter optimization, we performed 4-fold cross-validation in the training process. As a result, these models were developed by applying 4-fold cross-validation to 80% of the total samples. Test sets comprising the remaining 20% of the whole samples were used to evaluate the performance of the models. All models were trained 20 times by different seeds to avoid obtaining model bias. Finally, the model performance was assessed based on the averaged results.

In the training processes of ANN models, the coefficient of determination (R^2^) and mean square error (MSE) were used to evaluate the accuracies and losses. After optimization of hyperparameters, we used an *Adam* optimizer with a batch size of 256, learning rate of 0.00001, drop-out of 0, and weight decay of 0.01 to train these models. To obtain the number of iterations when the model was optimal, we repeatedly tested the variation of R^2^ and MSE with the number of iterations (Fig. S1). The results showed that after reaching 5000 iterations, the R^2^ and MSE of most models started to remain stable. The number of iterations for all models was set to 10000, considering the trade-off between the time cost of iteration and the lowest losses.

### From neutral community model to neutral and non-neutral partitions

To determine the potential importance of stochastic processes on WWTP community assembly, we used a neutral community model (NCM) to predict the relationship between an ASVs’ occurrence frequency and their relative abundance across the wider metacommunity(28). The model is an adaptation of the neutral theory adjusted to large microbial populations. In this model, m is an estimate of dispersal between communities, being the estimated migration rate. Because the estimation of migration rate m is affected by the number of sequences in samples, we flattened the number of sequences in each sample to 2000 before fitting the neutral community model, allowing us to compare estimated migration rates for different microbial partitions.

In this study, all 46628 ASVs were separated into three partitions depending on whether they occurred more frequently than (above partition), less frequently than (below partition), or within (neutral partition) the 95% confidence interval of the NCM predictions(29). To explore the effect of the potential migration rate of ASVs on their predictability, we analyzed and compared the predictive accuracy of different (above, neutral, and below) partitions belonging to the common ASVs_>10%_ subcommunity.

### Definition of core taxa

A global-scale core microbial subcommunity of WWTP was determined based on multiple reported measures. In this report, we explored the predictability of microbial taxa at the ASV level (100% similarity), as such the classification criteria for core ASVs were slightly different than those for core OTUs(3). First, ‘overall abundant ASVs’ were filtered out according to the mean relative abundance (MRA) across all samples. We selected all top 1% ASVs as overall abundant ASVs. Second, ‘ubiquitous ASVs’ were defined as ASVs with an occurrence frequency in more than 20% of all samples. Finally, ‘frequently abundant ASVs’ were selected based on their relative abundances within a sample. In each sample, the ASVs were defined as abundant when they had a higher relative abundance than other ASVs and made up the top 80% of the reads in the sample(30). A frequently abundant ASV was defined as abundant in at least 10% of the samples. Following the same criteria described above, a core ASV should be one that was from the top 1% ASVs, a core ASV also had to be detected in more than 20% of the samples and dominant for more than 10% of the samples. Corresponding to the core taxa, ASVs that did not meet the above three conditions were called non-core taxa.

### Statistical analysis

All alpha diversity measures were conducted using the vegan and Picante packages of R (v. 4.0.3). Unless indicated otherwise, an unpaired, two-tailed, two-sample Student’s *t*-test was performed for comparative statistics using the *t*.*test* function in the stats package of R 4.0.3. Linear correlation analyses between different parameters were implemented using the *lm* function in the stats package of R 4.0.3. All analysis and graphing were done using R4.0.3 or python 3.8. The source code is available on https://github.com/Neina-0830/WWTP_community_prediction.

## Results

### Overview of microbial community structure in AS systems

By preprocessing 777 (no data leakage) activated sludge samples from 269 wastewater treatment plants located in 23 countries across 6 continents using the QIIME2 pipeline (see Methods for details), we obtained the basic information of microbial community structure in AS systems. Specifically, the Shannon-Wiener index ranged from 2.90 to 6.41(Fig. 1a), Pielou’s evenness index ranged from 0.50 to 0.90(Fig. 1b), species richness ranged from 217 to 2014(Fig. 1c), and Faith’s phylogenetic diversity ranged from 22.58 to 148.33(Fig. 1d). Detailed information about alpha diversity is provided in Table S3.

**Fig. 1.**
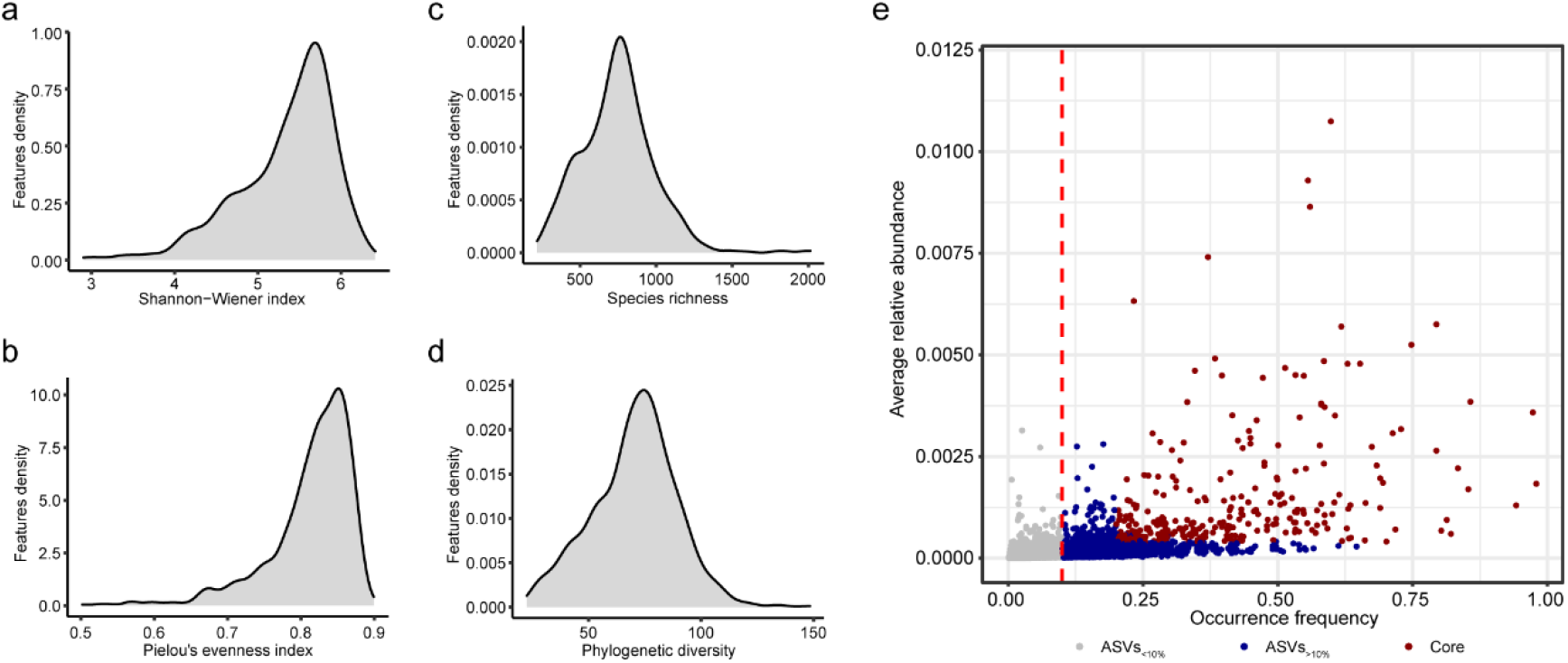
Overview of microbial community structure in AS system. Distribution of Shannon-Wiener index (**a**), Pielou’s evenness index (**b**), species richness (**c**), and Faith’s phylogenetic diversity (**d**). **e**. Occurrence frequency and average relative abundance distribution of all ASVs in the AS system.

In addition, we analyzed the distribution features of the average relative abundance and occurrence frequency of the ASVs. The results showed that ASVs in the AS systems were dominated by low relative abundance (Fig. 1e), which is in line with the general ecological environment(31). In this study, we predicted that only 1493 ASVs appeared in at least 10% of samples, of which 290 belonged to the core ASVs (Fig. 1e). We defined overall abundant, ubiquitous and frequently abundant ASVs as the core ASVs (see Methods for more details).

### Alpha-diversities of AS systems can be predicted by ANN models

#### Predictability of alpha-diversities

To obtain an overall prediction of AS community structure, we first constructed predictive models for different alpha diversity indices, including the Shannon-Wiener index, Pielou’s evenness index, species richness, and Faith’s phylogenetic diversity. Here, the predictive accuracy is measured relative to the 1:1 observed-predicted line (rather than a best-fit line), named R^2^_1:1_, so accuracy assessments are both qualitative and quantitative (18). By comparing the observed and predicted alpha diversities in test sets, we found that predictive accuracies R^2^_1:1_ of the Shannon-Wiener index (Fig. 2b), Pielou’s evenness index (Fig. 2c), species richness (Fig. 2d), and Faith’s phylogenetic diversity (Fig. 2e) were 60.42%, 54.11%, 49.92%, and 60.37%, respectively.

**Fig. 2.**
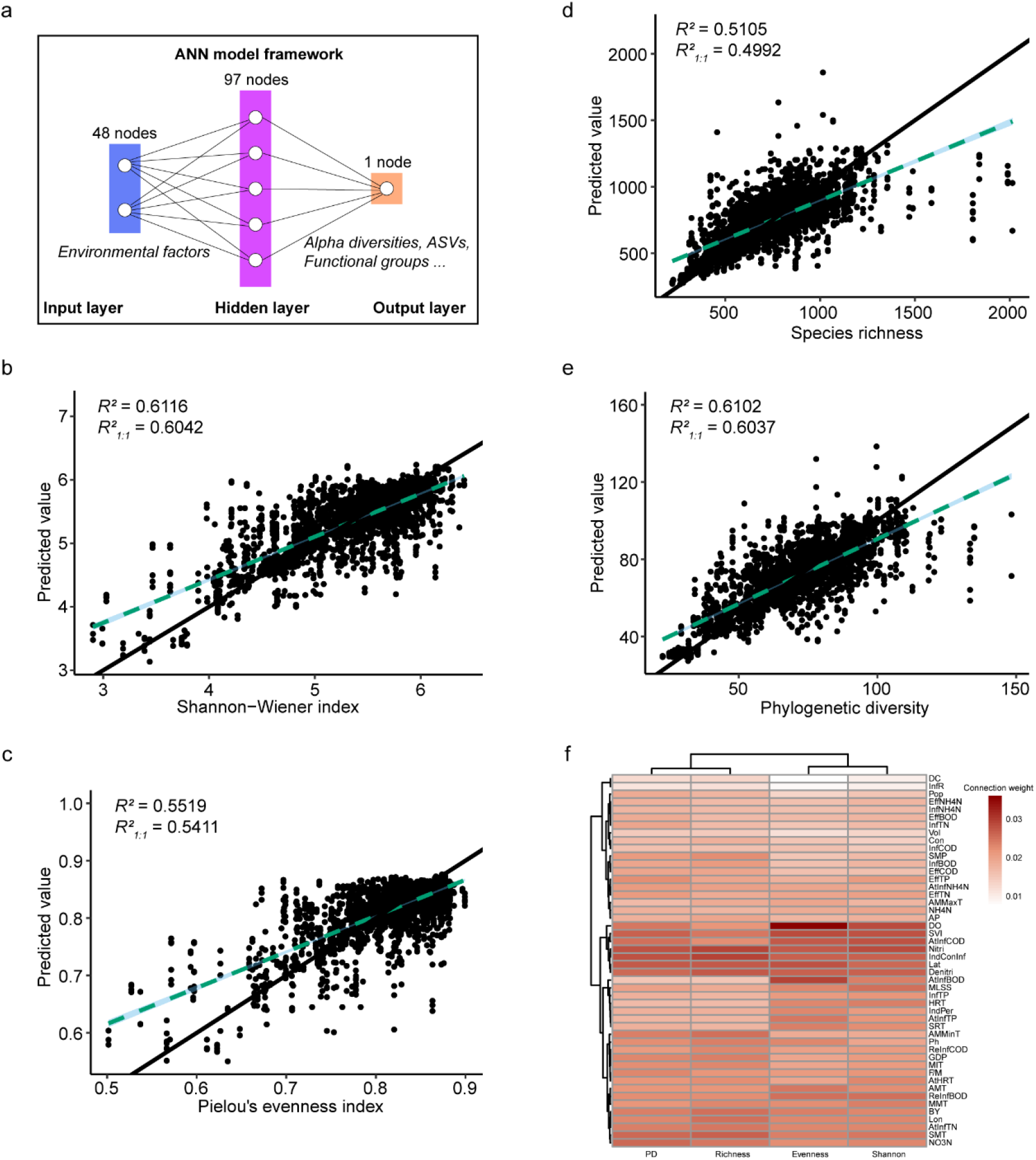
Prediction of alpha diversity. **a.** The framework of ANN models: input data (blue), output data (red), and a predictive model trained to compute output data from input data (purple). Correlations between observed and predicted values of Shannon-Wiener index (**b**), Pielou’e evenness index (**c**), species richness (**d**), and Faith’s phylogenetic diversity (**e**). The 1:1 relationship is shown as a solid black line, and the best fit is shown as the dashed light blue line. The blue shaded region represents the 95% confidence interval for the best fit line. We reported the R^2^ value of the best fit line between predicted and observed and the R^2^ observations relative to the 1:1 line. **f**. Heatmap of importance weights of environmental factors in alpha diversity predictive models.

Comparing the predictability of different alpha diversity indices, we found that the Shannon-Wiener index and Pielou’s evenness index were more predictable than species richness, which may be related to the environmental sensitivity of species evenness. Species evenness has previously been reported to be more sensitive to human activity and environmental changes than richness because environmental conditions may significantly affect ecosystems long before a species is threatened by extinction(32). In addition, the predictive accuracy of phylogenetic diversity was also higher than species richness, reflecting that species’ evolutionary history may be influenced by environmental factors surrounding the microbial community.

#### Environmental factors important for predicting alpha-diversities

During the model training process for predicting alpha-diversities of AS systems, an importance weight value was assigned to each environmental factor by the Garson’s connection weight method(33). The factors with higher importance weights were more informative when the model was used to predict alpha diversities.

To assess the importance of different environmental factors in predicting alpha-diversities of AS microbial communities, we ranked the average importance weights of environmental factors in different predictive models in descending order (Fig. S2). The results showed that the physicochemical property DO was most important for predicting the Shannon-Wiener and Pielou’s evenness indices, but inflow related industry wastewater source (IndConInf) was most important for predicting species richness and Faith’s phylogenetic diversity. Climatic condition Lat, design related N removal processes (Nitri and Denitri), inflow condition AtInfCOD, and the physicochemical parameter SVI were also environmental factors with high average importance weights for predicting alpha diversities (Fig. 2f).

### Assessment of the predictivity of community structure using ANN model

#### Predictability of the relative abundances of ASVs

To obtain a deep prediction of AS community structure, we predicted the relative abundance of ASVs in AS systems. We constructed predictive models for the 1493 ASVs found in more than 10% of samples (ASVs_>10%_) (see Methods for details), which accounted for 3.2% of the total ASVs and 64.97 ± 0.54% (mean ± SEM) of the relative abundance in AS samples (Fig. 3a). The results showed that the average predictive accuracy R^2^_1:1_ of ASVs_>10%_ was 35.09% (Table S4). Further, we found that 19.83% of ASVs_>10%_ could be predicted with R^2^_1:1_ over 50%, 60.82% of ASVs_>10%_ could be predicted with R^2^_1:1_ over 30%, and 91.96% of ASVs_>10%_ could be predicted with R^2^_1:1_ over 10% (Fig. 3b).

**Fig. 3.**
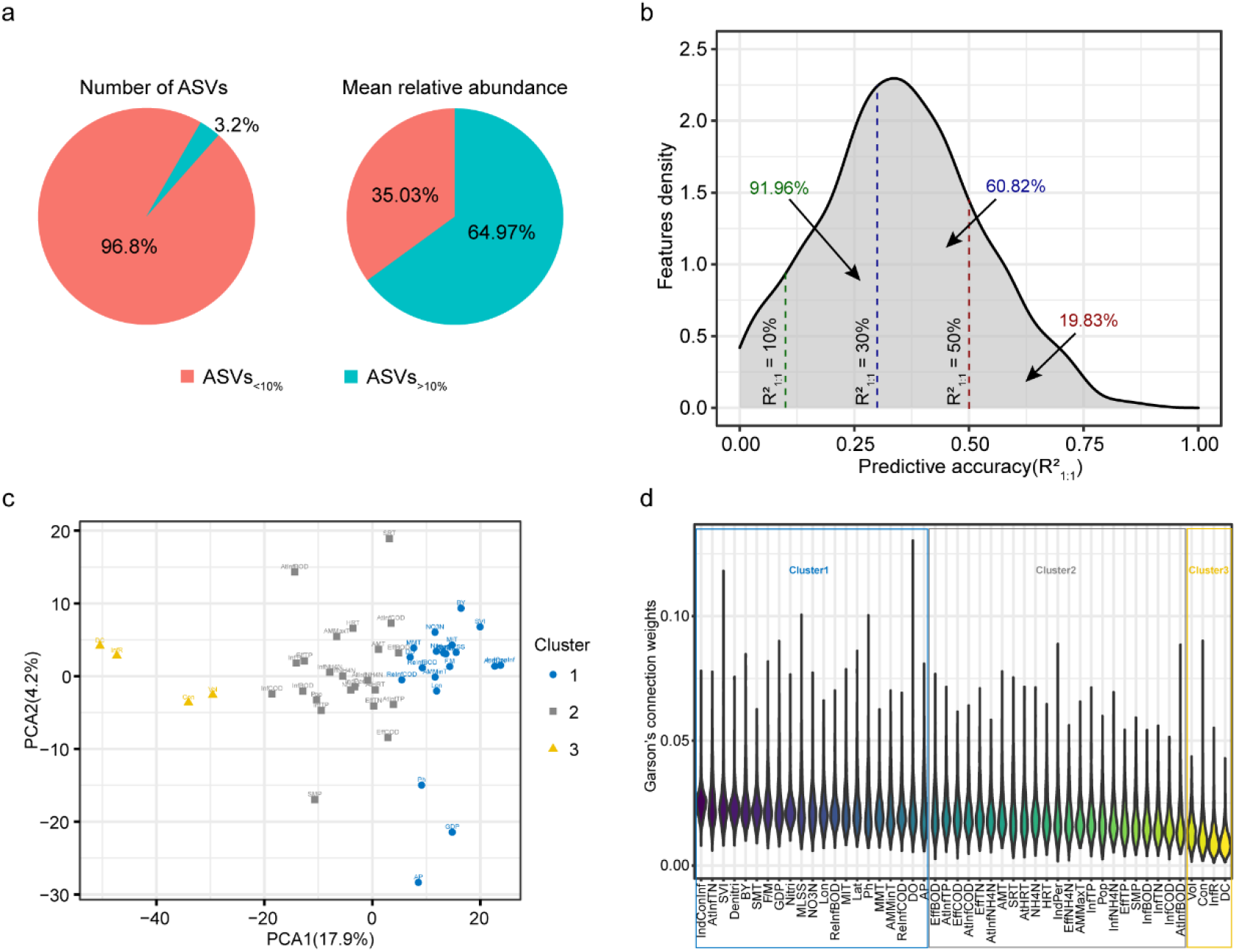
Prediction of the relative abundance of ASVs_>10%_. **a**. Percentage of ASV number and relative abundance of ASVs_<10%_ versus ASVs_>10%_. **b**. Distribution of the predictive accuracy of ASVs_>10%_. The dark green, dark blue and dark red text represent the proportion of ASVs with prediction accuracy exceeding 10%, 30% and 50%, respectively. **c**. Principal component analysis (PCA) of environmental factors colored by k-means clusters. **d**. Ranking of environmental factors in descending order of median importance weights.

In addition, we also predicted the structures of the microbial communities of the test samples, by recovering the ASVs_>10%_ subcommunity of each sample in its entirety (21). Here, we refer to the observed values of ASVs_>10%_ subcommunities in different test samples as “observed communities”, and the corresponding predicted values as “predicted communities”. By comparing the intra-group and inter-group differences between the predicted and observed communities, we found that the Bray-Curtis similarity between intra-groups was significantly higher than that between inter-groups (Fig. S3a). This result also proved that the ANN model could predict the microbial community structure of the AS system from an overall perspective.

Furthermore, we predicted microbial taxa at different taxonomic levels and found that microbial community structure had high average predictive accuracy ranging from 33.32% to 41.6% at all taxonomic levels (Fig. S3b). The predictive accuracy R^2^_1:1_ of the three most abundant phyla (Proteobacteria, Bacteroidota and Myxococcota) in the AS system were 64.54%, 55.37% and 59.04%, respectively. The three most abundant orders Burkholderiales, Chitinophagales and Pseudomonadales could be predicted with R^2^ of 63.89%, 56.19%, and 42.81%, respectively (Table S5).

#### Importance of environmental factors in the prediction of ASVs

During the model training process for predicting abundances of ASVs, an importance weight value was also assigned to each environmental factor as above (Table S6). By displaying the importance weights of environmental factors in different ASVs predictive models in descending order of their mean values, we found that environmental factors had different weights in predicting different ASVs (Fig. S4a). Further, we clustered the environmental factors into three clusters according to their importance weights using k-means clustering algorithm and displayed them using principal components analysis (PCA) (Fig. 3c). We found that these three clusters corresponded to three parts divided by the median of importance weights in descending order (Fig. 3d). This result showed that environmental factors of cluster1, which included climatic condition SMT, design and operation parameters BY and Denitri, inflow conditions IndConInf and AtInfTN, and physicochemical properties SVI etc, contributed the most to the prediction of community structure, cluster2 was second, and cluster3 was the least important group of factors in predicting community structure (Table S7).

We then wondered what influences the importance weights of environmental factors in predicting microbial taxa. Before constructing the predictive model, we performed a Mantel test analysis on the correlation between the ASVs_>10%_ subcommunity and ecological environment factors (Fig. S4b). The correlation analysis showed that environmental factors significantly associated with the ASVs_>10%_ subcommunity included climate conditions Lat, Lon, AMT, AMMinT, SMP, and GDP, design and operation parameter Nitri, and physicochemical property MIT (Table S7; Pearson’s ρ>0.2, p<0.01). By comparing the importance of environmental factors in predicting community structure and the correlation between environmental factors and community structure, we found that some of the environmental factors that had high importance weights in many predictive models were not strongly correlated with the ASVs_>10%_ subcommunity (Fig. S5). For example, inflow conditions IndConInf and AtInfTN, which were important for predicting the relative abundance of ASVs, did not significantly correlate with the ASVs_>10_ subcommunity. However, despite these differences, there was a significant positive correlation between the importance weights of environmental factors and their correlation coefficients with the ASVs_>10%_ subcommunity (Fig. S6a, R^2^=0.1271, p<0.05). These results showed that in addition to the correlation analysis, importance weights analysis in ANN predictive models also helped to expand the range of environmental factors that should be paid attention to when exploring the performance of WWTPs.

In addition, we also analyzed the influence of the distribution of environmental factors on their weights. The result showed that both the skewness (Fig. S6b; R^2^=0.6268, p<0.001) and kurtosis (Fig. S6c; R^2^=0.7106, p<0.001) of normalized environmental factors were significantly negatively correlated with their average importance weights in predicting ASVs. This suggested that environmental factors of low skewness and low kurtosis may be more important in predicting community structure.

### Characteristics of ASVs with high predictabilities

#### ASVs with higher relative abundances and occurrence frequencies can be better predicted using the ANN model

To investigate correlation between the predictability of ASVs and their distribution features, we compared the predictability of ASVs with different relative abundances and frequencies. The correlation analysis between the predictive accuracy R^2^ of all 1493 ASVs_>10%_ and their relative abundances showed that the R^2^_1:1_ of an ASV was significantly positively correlated with its relative abundance (Fig. 4a; R^2^=0.05279, P<0.001), indicating that the high predictability of an ASV may be related to its high relative abundance. Furthermore, we found that the R^2^_1:1_ of an ASV was slightly positively associated with its occurrence frequency at significant levels (Fig. 4b; R^2^=0.02602, P<0.001), indicating that the high predictability of abundant ASV_>10%_ may also be related to its high occurrence frequency. Further, we grouped ASVs based their relative abundances and occurrence frequencies as in previous studies (see more details in Supplementary information S1), and the results showed that ASVs_>10%_ with high relative abundance and high occurrence frequency were significantly more predictable than those with low relative abundance (Fig. S7a) and low occurrence frequency (Fig. S7b), which is consistent with the previous result. It is worth mentioning that the occurrence frequency of an ASV has a significant positive correlation with its relative abundance (Fig. S7c, R^2^=0.2978, P<0.001), supporting that high relative abundance and high occurrence frequency can corroborate each other in their contribution to predictability.

**Fig. 4.**
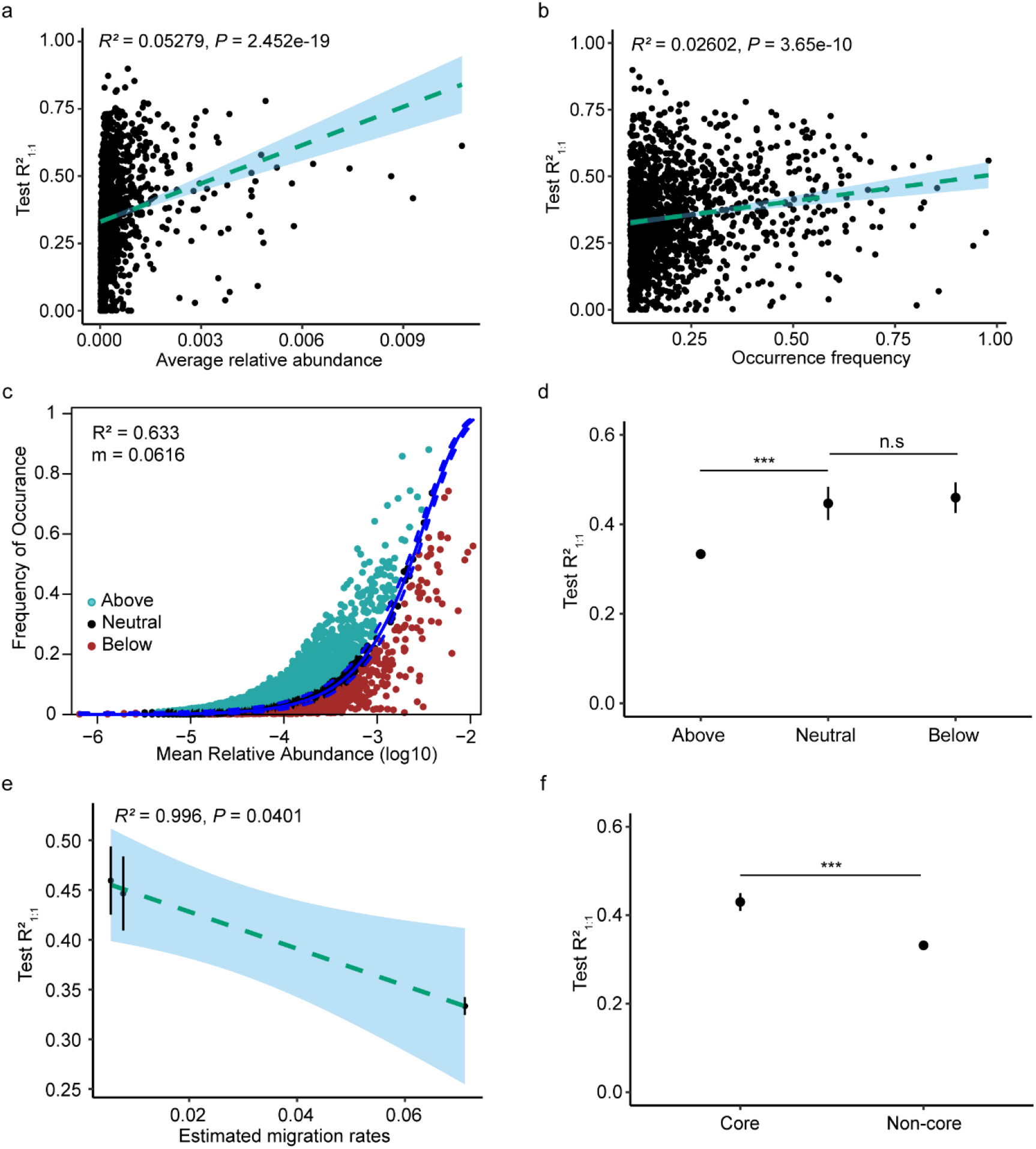
Distribution features and predictability of ASV_>10%_. **a.** Correlation of test R^2^_1:1_ with the average relative abundance. **b**. Correlation of test R^2^_1:1_ with the occurrence frequency. **c**. Fit of the neutral community model (NCM) of AS system community assembly. The solid blue lines indicate the best fit to the NCM as in Sloan et al. [31], and the dashed blue lines represent 95% confidence intervals around the model prediction. R^2^ indicates the fit to this model, and m indicates the estimated migration rate. **d**. The test R^2^_1:1_ of above, neutral, and below partitions. **e**. Correlation of test R^2^_1:1_ with the estimated migration rate. The data was provided by the results of different partitions. **f**. The test R^2^_1:1_ of core and non-core taxa. Statistical analysis was performed using a two-sample Student’s *t*-test: ***, p < 0.001; n.s, p > 0.05, no significance.

Previous studies had demonstrated that rare taxa were more dynamic than abundant taxa(34), so we wondered whether a taxon’s predictability was related to its variability across samples. To explore this question, we analyzed the relationship between R^2^_1:1_ of the ASVs and their coefficients of variation, and found that the predictive accuracy of an ASV was significantly negatively correlated with its coefficient of variation (Fig. S7d; R^2^= 0.01946, P<0.001), implying that taxa with higher variability were less predictable.

#### The predictability of an ASV decreases as its potential migration rate increases

Previous studies have demonstrated that the process of community assembly is closely related to its predictability(18, 35, 36), so we explored the association between microbial community assembly mechanisms in AS systems and the predictability of the corresponding taxa. By neutral community model (NCM) model fitting, we found that stochastic processes explained 63.3% of the microbial community variation in AS systems (Fig. 4c). The ASVs_>10%_ were subsequently separated into three partitions (above, below and neutral) depending on their occurrence frequencies and relative abundance (see Methods for details). By comparing distribution features of the three partitions, we found that the relative abundance (Fig. S8a) and occurrence frequency (Fig. S8b) of ASVs_>10%_ in the below partition were significantly higher than those of the above partition. Further, we found that the predictive accuracy R^2^_1:1_ of the below partition was also significantly higher than that of the above partition (Fig. 4d). This result again showed that ASVs with higher relative abundances and occurrence frequencies can be better predicted using ANN models.

In addition, the estimated migration rates of the different partitions assessed by NCM were also different. Points above the fitting curve represent taxa found more frequently than expected, suggesting that they have a higher migration ability and can disperse to more locations. Points below the fitting curve represent taxa found less frequently than expected, suggesting their lower dispersal ability in WWTPs on a global scale or that they have a stronger response to local environmental conditions. The fitting results also confirmed that the taxa in the above partition had the highest estimated migration rates, and the taxa in the below partition had the lowest estimated migration rates (Fig. S9). Further analysis of the relationship between the migration rate and predictability of different partitions, we found that a taxon’s predictability had a high negative correlation with its estimated migration rate (Fig. 4e, R^2^=0.996, P=0.0401), indicating that the predictability of an ASV decreased as its potential migration rate increased.

#### Core taxa of AS systems can be predicted by ANN models

Due to its highly complex organic environment, activated sludge selects a unique core community that does not overlap with the core communities of other habitats(3). We evaluated the predictabilities of core taxa that were abundant and ubiquitous (see Methods for details) using ANN models. As defined in Methods, we obtained 290 core ASVs and 1203 non-core ASVs in the ASVs_>10%_ subcommunity (Fig. S10a). Our analyses found that the relative abundance (Fig. S10b) and occurrence frequency (Fig. S10c) of core taxa were significantly higher than those of non-core taxa, and the estimated migration rate of core taxa was lower than that of non-core taxa (Fig. S10d).

By assessing the predictability of the core taxa, we found more than 37.59% of the core ASVs could be well predicted with an R^2^ of over 50%, more than 78.62% could be well predicted with an R^2^ of over 30%, and more than 94.48% could be well predicted with an R^2^ of over 10%, and the average prediction accuracy was 42.99% (Table S4), which was significantly higher than the non-core taxa (Fig. 4f). Because the core taxa are reported to be closely related to nitrogen removal, phosphorus removal, and even flocculation enhancement of activated sludge(12, 30, 37), the results implied that the ANN model could be used to assess the performance of WWTPs by predicting the dynamics of the core taxa.

### Prediction of major functional groups in AS systems

To more directly understand and control the performance of WWTPs, we predicted and analyzed major functional groups of microbial communities in the AS system using ANN models. Referring to the MiDAS4 database, the functional groups in AS systems include nitrogen removal groups (nitrifiers and denitrifiers), phosphorus removal groups (PAOs), and their competitors (GAOs), and filamentous organisms(26) (Table S8). Then, we calculated the total relative abundance of different functional groups in each sample by summarizing the relative abundances of ASVs from the same functional groups. Finally, we predicted the total relative abundance of each functional group using the ANN model. The results showed that the predictive accuracy R^2^_1:1_ for nitrifiers, denitrifiers, PAOs, GAOs, and filamentous organisms was 32.62%, 62.65%, 53.46%, 53.31%, and 62.86%, respectively (Fig. 5a, Table S9).

**Fig. 5.**
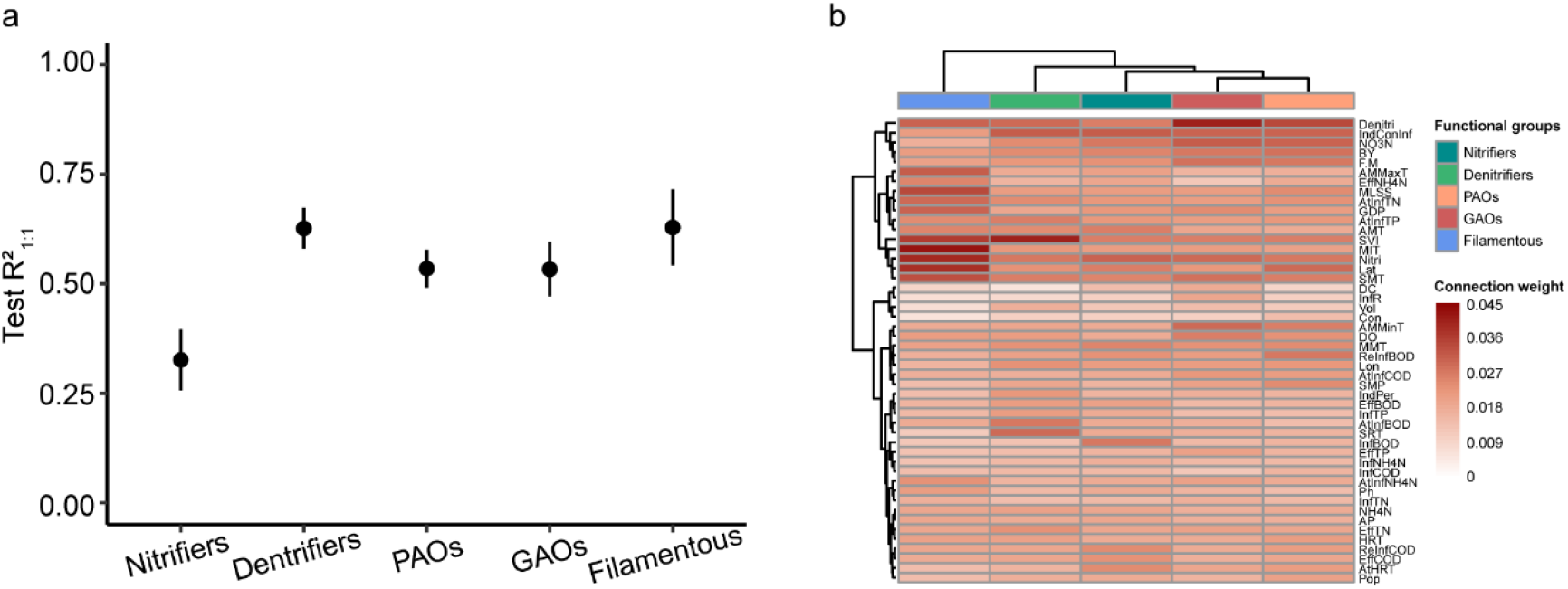
Prediction of functional groups. **a.** The test R^2^_1:1_ of nitrifiers, denitrifiers, PAOs, GAOs, and filamentous organisms. **b**. Heatmap of importance weights of environmental factors in the predictive models of functional groups. Errors bars in these graphs show the 95% credible intervals of the mean values. Statistical analysis was performed using a two-sample Student’s *t*-test: ***, p < 0.001; n.s, p > 0.05, no significance.

To further understand the prediction results of functional groups, we also analyzed the importance weights of environmental factors in their predictive models. By performing Ward clustering analysis on the importance weights of environmental factors in these predictive models of different functional groups, we found that the importance of environmental factors in predicting PAOs and GAOs was the most similar, followed by nitrifiers and denitrifiers (Fig. 5b), which implied a consistent contribution of environmental factors when predicting relevant functions. Overall, the design and operation parameters BY and Denitri, inflow condition IndConInf, and physicochemical properties F/M and NO3N were important for predicting nitrogen and phosphorus removal function, while climatic conditions Lat and SMT, design and operation parameter Nitri, and physicochemical properties SVI and MIT significantly affected the prediction of filamentous organisms. Additionally, SVI also had a crucial impact on the prediction of denitrifiers, which may be because some filamentous bacteria also function as denitrifiers(38). To demonstrate the importance of these environmental factors, we only used the above 10 high-weight environmental factors to predict functional groups. The results showed that using only those ten factors allowed us to predict the abundance of major functional groups in AS systems with a predictive accuracy of R^2^_1:1_ ranging from 17.25% to 52.00% (Fig. S11).

In summary, the climatic conditions Lat and SMT, the design and operation parameters BY, Denitri and Nitri, the inflow condition IndConInf, as well as some sample physicochemical properties (F/M, SVI, MIT and NO3N) of AS systems all affect the prediction of functional groups. Controlling these critical environmental factors can help us regulate the performance of WWTPs, which will guide us to design more reasonable operating parameters according to environmental changes.

## Discussion

In this study, we predicted the diversity and the structures of the microbial community, as well as the functional groups in AS systems using ANN models. We also evaluated the importance of environmental factors in the predictions.

The use of artificial neural network (ANN) models in this study increased the predictive power of complex systems of microbial communities. When ANN models were used to predict ASVs appearing in at least 10% of the samples, 60.82% of the ASVs_>10%_ had a prediction accuracy R^2^_1:1_ exceeding 30%. In a previous study, the multiple regression model could only explain about 15% of the variability in genus-level taxonomy of a soil bacterial microbial community(18) and only predicted the top ten taxa of that community. Compared with this previous study, our prediction accuracy was greatly improved, with prediction range being increased to all ASVs appearing in at least 10% of samples, which proves the application potential of ANN models in predicting the complex systems of microbial communities.

Using the Neutral Community Model (NCM) proposed by Sloan et al.(28), this study transformed migration from a vague qualitative concept into a number with biological meaning, the potential migration rate (m). Higher values of m indicate that a species is less limited by dispersal. The low migration rate of high-abundance taxa and high migration rate of low-abundance taxa in this study (Fig. S12) indicates that dispersal limitation has a significant effect on high-abundance taxa, but not on low-abundance taxa, which is consistent with findings for some ecosystems(39, 40). High-abundance taxa with a low migration rate will appear in some samples due to environmental selection(41), and their relative abundance can be well predicted using these environmental factors. However, low-abundance taxa with high migration rates usually appear in a sample when the migration occurs and the spatial heterogeneity of the sample provides them with ecological niches. Neither the randomness of migration nor the spatial heterogeneity of samples was reflected in our input environmental variables, as such, these environmental factors were less predictive of low-abundance taxa. In addition, low-abundance taxa have been reported to have higher abundance variability than high-abundance taxa(34), and prediction targets with higher variability are not conducive to the stability of the predictive model, further explaining why the predictability of the relative abundance of high-abundance taxa was significantly higher than that of low-abundance taxa.

The weight of environmental factors in the predictive model reflects the influence of environmental factors on the corresponding prediction targets to a certain extent. For example, our results showed that the most important environmental factors affecting the prediction of evenness and richness were DO and IndConInf, respectively. Evenness and richness are two critical indicators to measure the diversity of ecological communities. The former describes species differences, and the latter describes the number of species. Previous studies have demonstrated that relative abundances of some functional taxa are sensitive to changes in DO(42, 43), and the abundance of these functional bacteria reflect the differences in species abundance of the community. Therefore, DO has a high weight in predicting the evenness of microbial communities in AS systems. Industrial wastewater contains many toxic and harmful substances(44, 45), in which many microorganisms cannot survive. Therefore, industrial wastewater directly affects the population of microorganisms(46), and IndConInf plays an important role in predicting the richness of microbial communities in AS systems. The environmental factors with top weights in predictive models of nitrogen removal-related taxa ASV6 and ASV142 were AtInfTN, Nitri, and NO3N (Table S6). This correspondence between functions and environmental factors indicates that environmental factors with high weights in predicting microbial taxa may play an essential role in environmental filtering in the deterministic process of community assembly.

Important factors that cannot be identified using traditional methods may be highlighted by ANN modeling. Conventional studies on AS systems have only focused on the correlation relationship between environmental factors and microbial communities(16, 17, 47), which limited the scope of consideration for key environmental factors. For example, some previous studies had shown apparent differences between the microbial communities of industrial sewage and municipal sewage(14, 48), which showed that the IndConInf variable might impact the AS system’s microbial community (49). However, despite its high importance weight in ASVs_>10%_ predictive models, IndConInf was not significantly associated with the microbial community structure (Table S7). By analyzing the importance weights of environmental factors in predictive models, this study illuminated variables that require further attention and that can better predict and control the microbial community of AS systems.

Although our work has made some contributions to the prediction and interpretation of the microbial community structure in AS systems, we still cannot explain the weights of some environmental factors in the predictive model due to the black-box characteristics of the ANN model. Our results show that environmental factors with low skewness and low kurtosis distribution are more likely to have higher weights in predicting the relative abundance of microbial taxa, which we cannot explain using current knowledge. Increasing the interpretability of the ANN model will help us better use this powerful predictive tool to analyze our concerns, which is also the future direction of machine learning-based big data analysis.

Our results demonstrate that ANN models can predict the microbial community of AS systems well, including alpha diversity, ASVs_>10%_, core taxa, and major functional groups of AS systems. In addition, high relative abundance, high occurrence frequency, and low estimated migration rate all contribute to the predictability of taxa. These findings shed light on the role of the ANN model in predicting the complex system of microbial communities and provide new insights into the influence of environmental factors and the predictability of taxa.

## Acknowledgments

The authors thank the Global Water Microbiome Consortium (GWMC) and all the people involved for providing samples and plant metadata. This work was supported by the National Key R&D Program of China (2018YFA0902100 and 2021YFA0910300), the National Natural Science Foundation of China (91951204, 32130004, 32161133023, and 32170113), and the High-performance Computing Platform of Peking University.

## Author contributions

X.L. conceived the study and performed all analysis and computation. X.L. and Y.N. co-wrote the paper, and X.L.W. revised it. All authors discussed the results and commented on the article.

