## Supplementary information for "Predicting microbial community compositions in wastewater treatment plants using artificial neural networks"

### For

**This PDF file includes:**

S1. Grouping and Comparison of $ASVs_{>10\%}$

S2. Supplementary Figures 1 to 12

S3. Supplementary Tables 1 to 9

### S1. Grouping and Comparison of $ASVs_{>10\%}$

For a more detailed analysis of the impact of relative abundance and occurrence frequency on the predictability of ASVs, we also grouped all ASVs by relative abundance(1) and occurrence frequency(2), and compared the predictability of different groups.

First, grouped by relative abundance, there were 321 ASVs with an average relative abundance above 0.05%(named high-abundance taxa), 172 ASVs with an average relative abundance below 0.005%(named low-abundance taxa), and 1000 ASVs with an average relative abundance between 0.05% and 0.005% (named medium-abundance taxa) in the ASVs>10% subcommunity (Table S4). By comparing the predictability of high, medium, and low abundance taxa, we found that test $R^2_{1:1}$ of medium-abundance taxa was significantly higher than that of low-abundance taxa and lower than that of high-abundance taxa (Fig. S7a). This result suggested that the higher the relative abundance of microbial taxa, the more predictable it is.

Then, we divided the $ASVs_{>10\%}$ subcommunity into high (appears in more than 50% of samples), medium (appears in 20% to 50% of samples), and low (appears in less than 20% of samples) frequency groups (Table S4). By comparing the predictability of the different groups described above, we found that the test $R^2_{1:1}$ of the low-frequency group was significantly lower than that of the medium-frequency group, even though the predictive accuracy $R^2_{1:1}$ of the medium-frequency group and the high-frequency group had no significant differences (Fig. S8b). This result shows that the increase in occurrence frequency may also lead to higher predictability of the taxa.

### S2. Supplementary Figures

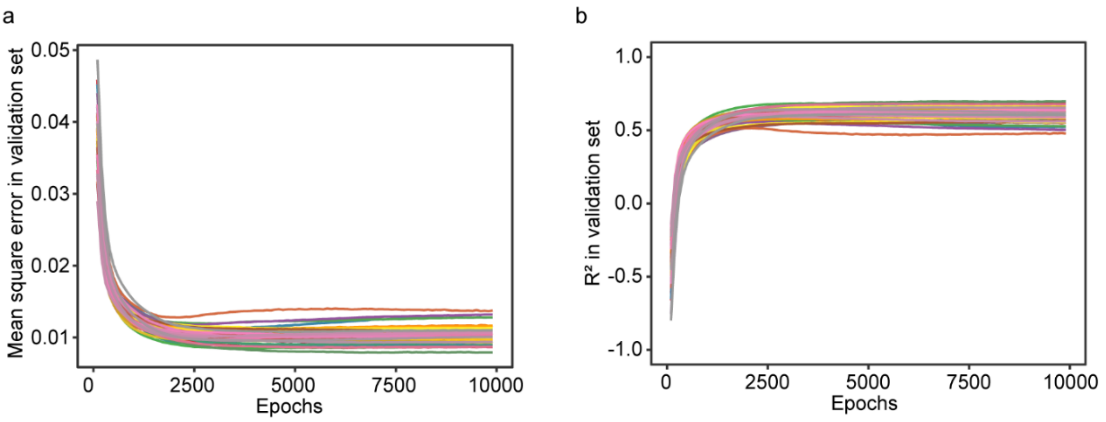


**Fig. S1** Changes of mean square errors (MSE) and coefficients of determination ($R^2$) on the validation set with epochs when training the model. Take the prediction of the Shanon-Wiener index as an example.

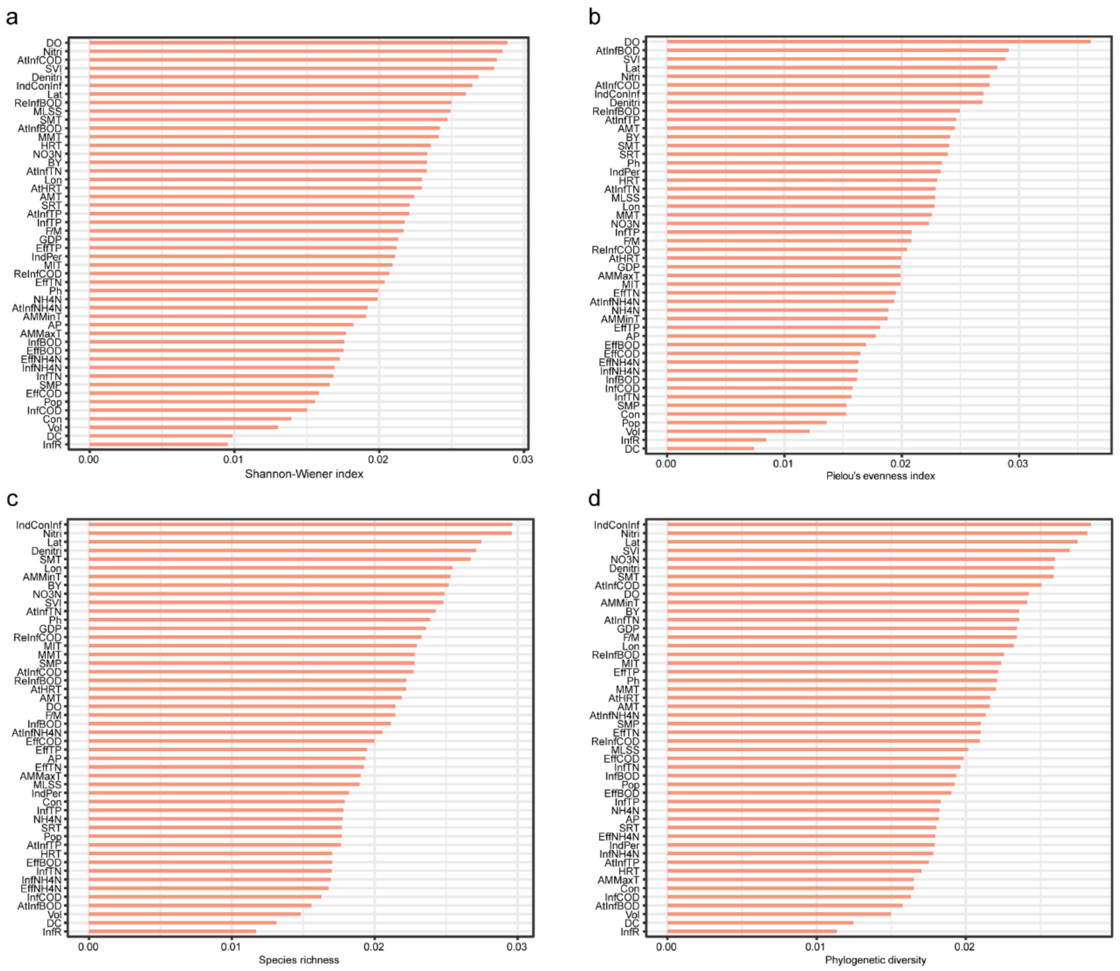


**Fig. S2** Ranking of importance weights of environmental factors in different alpha-diversities predictive models.

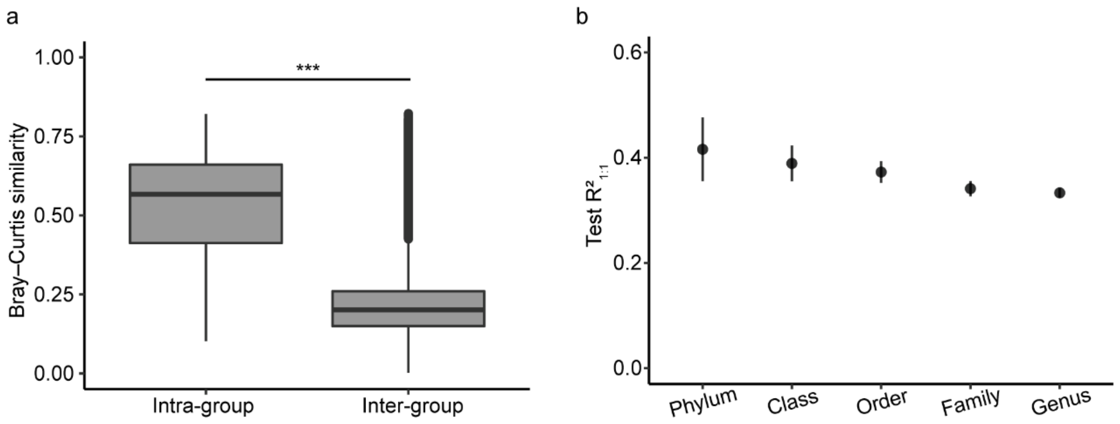


**Fig. S3 a.** Comparison of intra- and inter-group Bray-Curtis similarity between predicted and observed communities. Statistical analysis was performed using a two-sample Student's t-test: ***, $p < 0.001$. **b.** Average prediction accuracy $R^2_{1:1}$ of microbial taxa at different taxonomic levels.

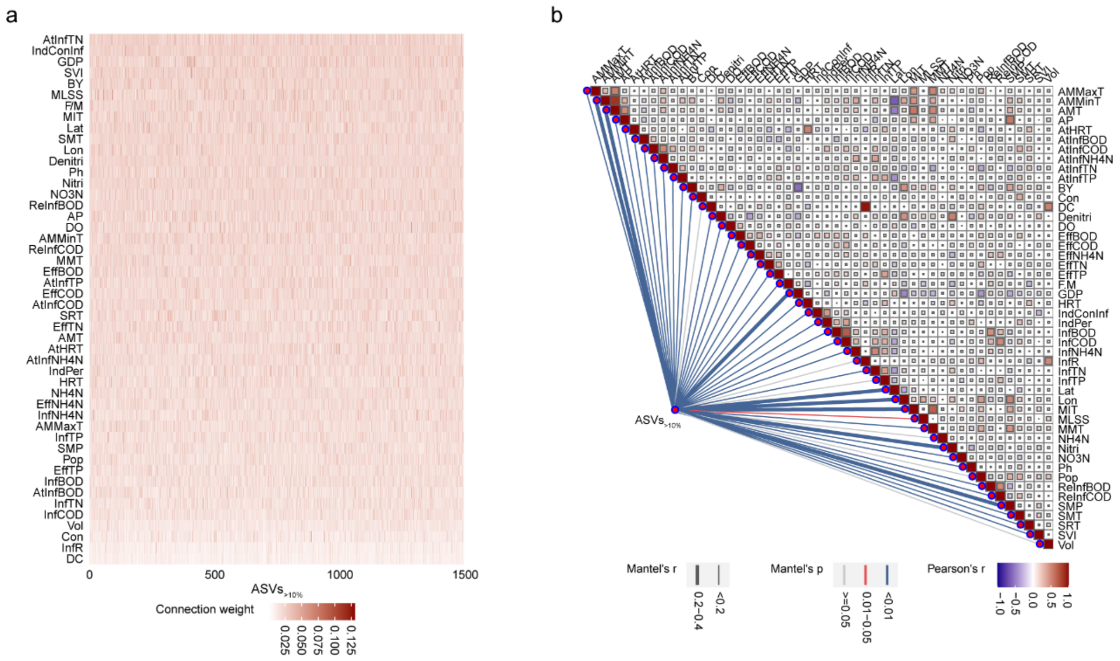


**Fig. S4 a.**Heatmap of importance weights of environmental factors in predictive models of ASVs belonging $ASVs_{>10\%}$ subcommunity. Environmental factors were sorted by average weights. **b.** Result of mantel tests on the correlation between $ASVs_{>10\%}$ subcommunity and ecological environment factors. Pairwise comparisons of environmental factors are shown with a color gradient denoting Pearson's correlation coefficient. The $ASVs_{>10\%}$ subcommunity (relative abundance of all ASVs belonging to $ASVs_{>10\%}$ subcommunity) of AS microbial community was related to each environmental factor by Mantel tests. Edge width corresponds to the Mantel's *r* statistic, and edge color denotes the statistical significance based on 999 permutations.

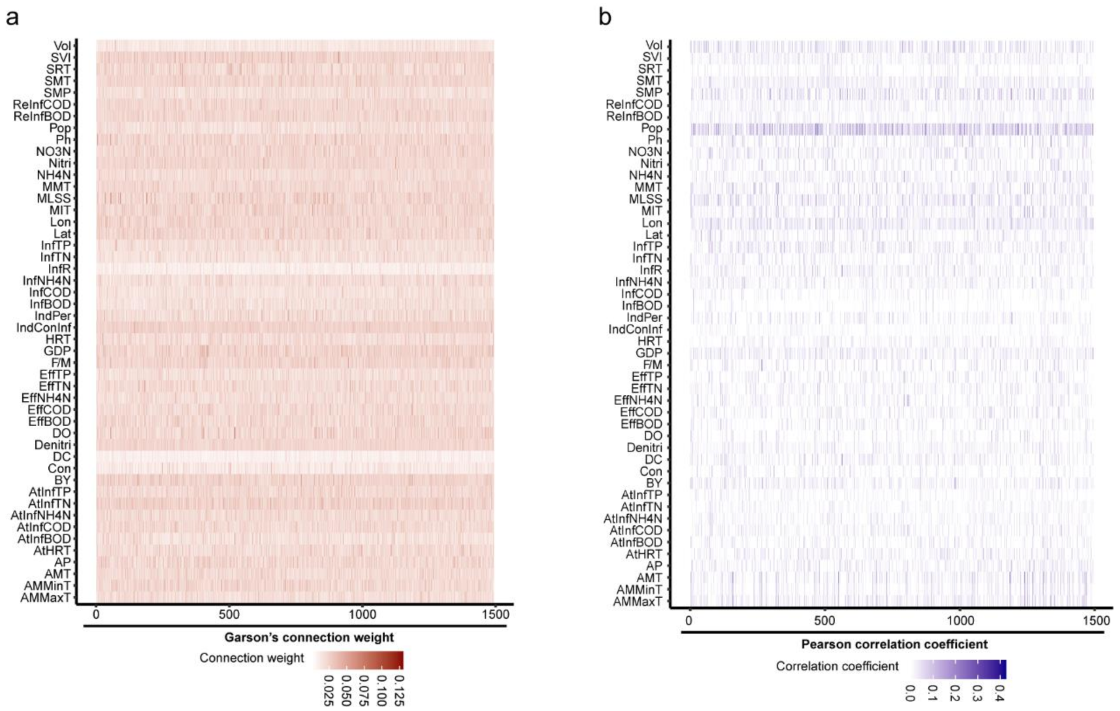


**Fig. S5 a.** Environmental factor importance weights for predicting ASVs$_{>10\%}$ subcommunity. **b.** Pearson's correlation coefficients between environmental factors and ASVs$_{>10\%}$ subcommunity.

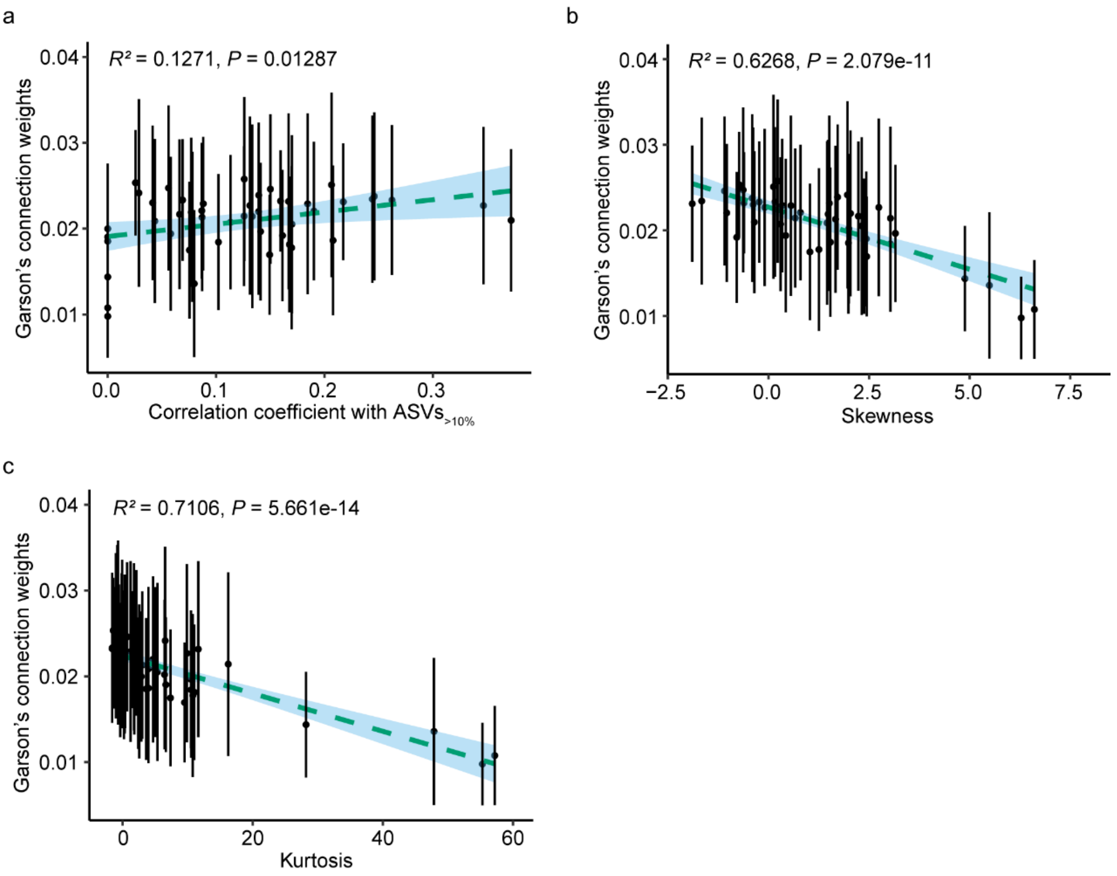


**Fig. S6 a.** Correlation of correlation coefficients of environment factors with ASVs$_{>10\%}$ subcommunity with their Garson's connection weights. Correlation of skewness (**b**) and kurtosis (**c**) of normalized environment variables with their Garson's connection weights. The best fit is shown in the ocean dashed line. The shaded sky region represents the 95% confidence interval for the best fit line. Points represent average Garson's connection weights, and errors bars show 95% credible intervals of average Garson's connection weights.

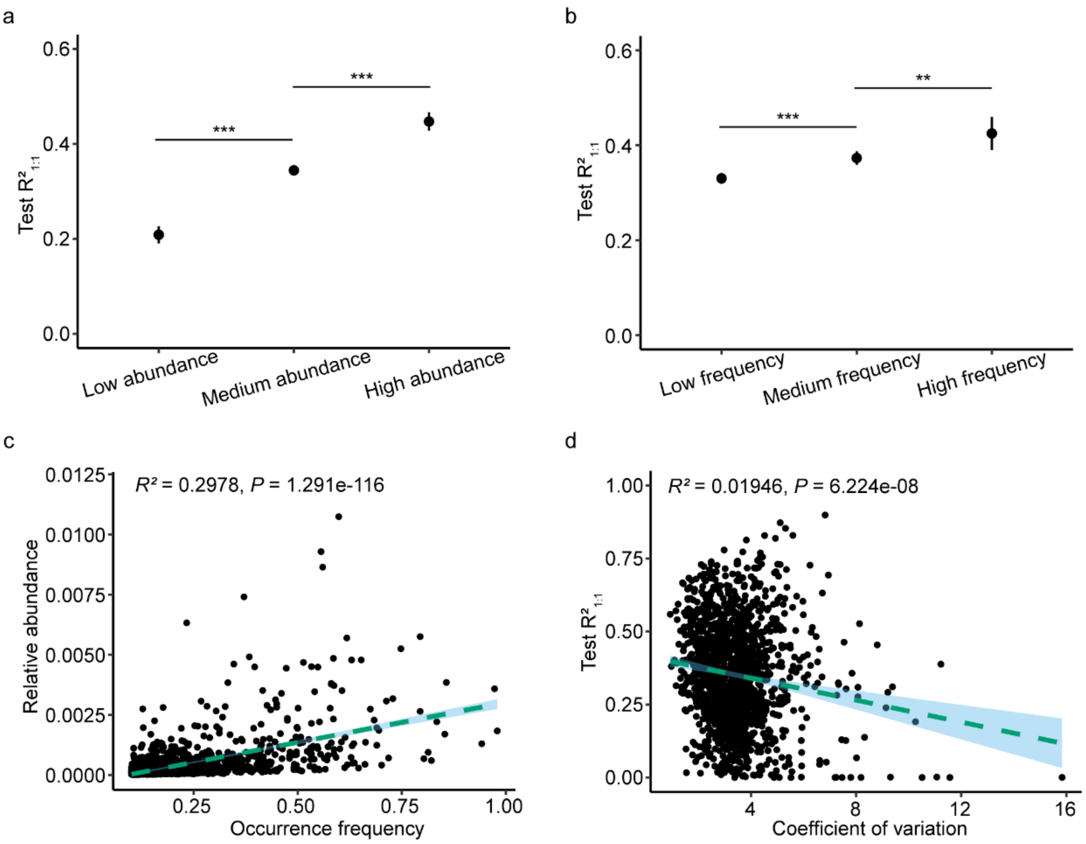


**Fig. S7 a.** Comparation of predictive accuracy $R^2_{1:1}$ between low, medium, and high abundance taxa. **b.** Comparison of predictive accuracy $R^2_{1:1}$ between low, medium, and high frequency taxa. **c.** Correlation of relative abundance with occurrence frequency of ASVs. **d.** Correlation of the $R^2_{1:1}$ in test sets with the coefficient of variation of ASVs. The date was provided by all ASVs in $ASVs_{>10\%}$ subcommunity. We reported the $R^2$ and P value of the best fit line in these figures.

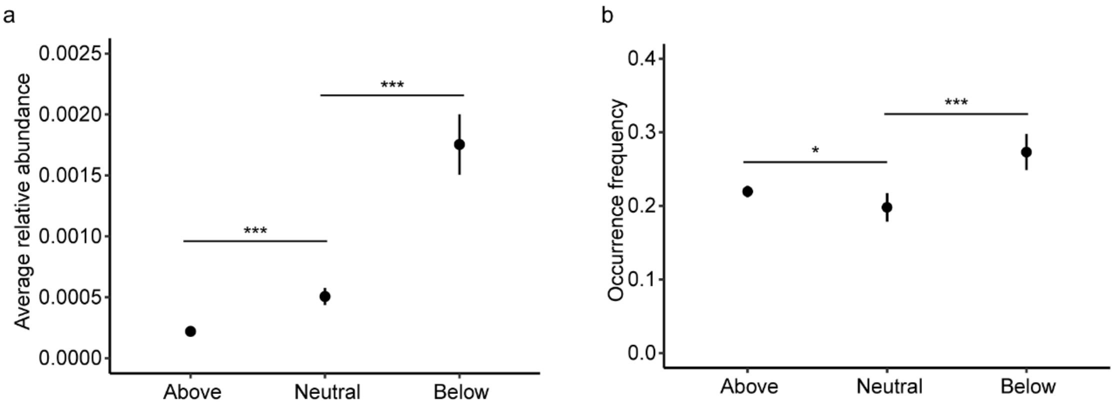


**Fig. S8** Comparation of average relative abundance (**a**) and occurrence frequency (**b**) between above, neutral, and below partitions.

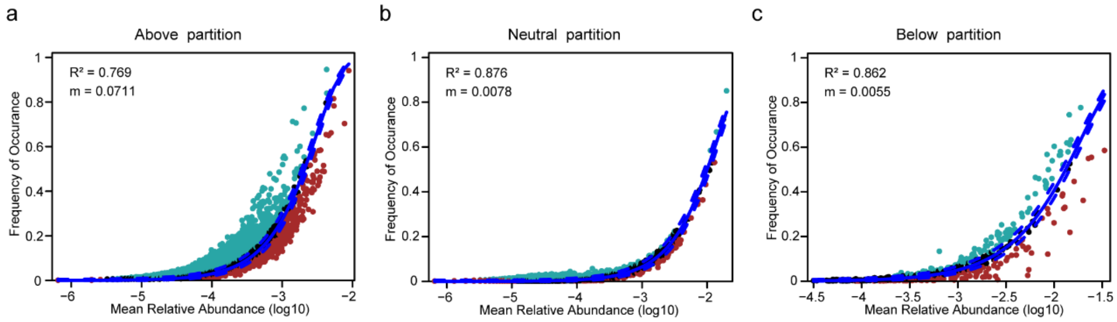


**Fig. S9** Fit of the neutral community model (NCM) of above (**a**), neutral (**b**), and below (**c**) partitions.

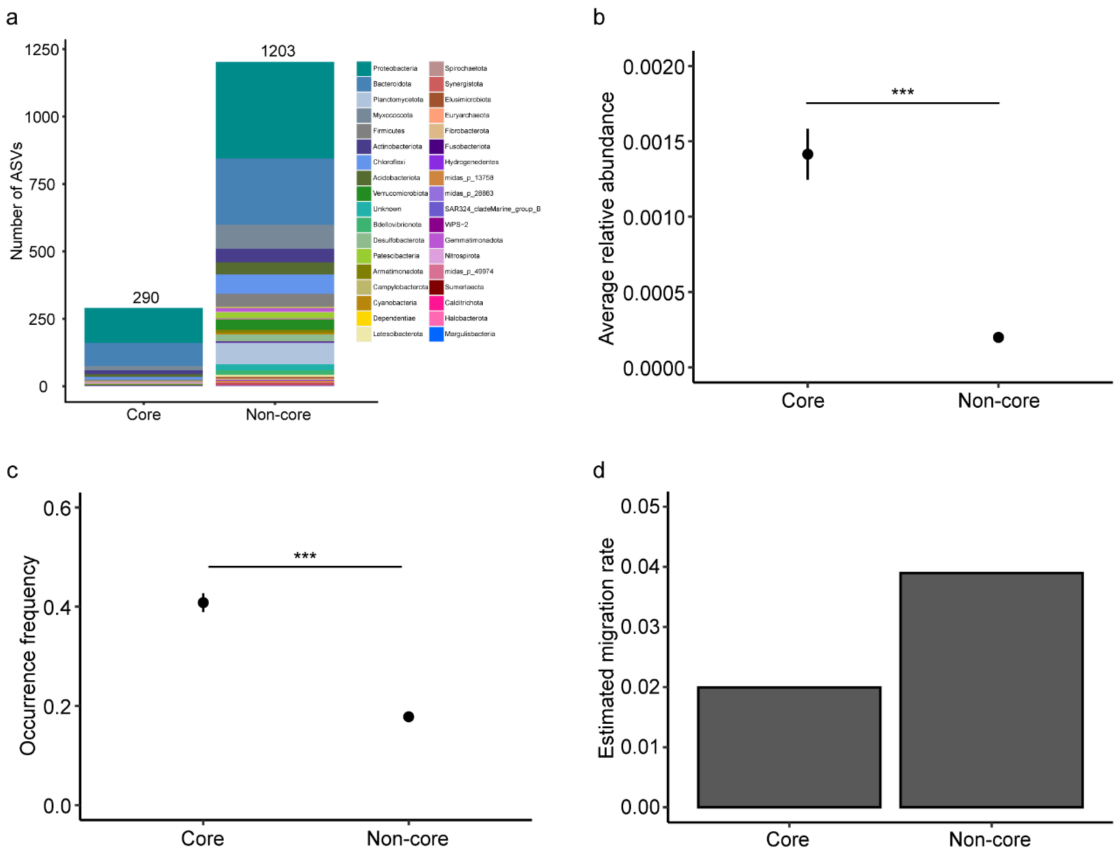


**Fig. S10 a.** The taxonomic composition of the core and non-core taxa at the phylum level. Comparation of average relative abundance (**b**) and occurrence frequency (**c**) between core and non-core taxa. **d.** The estimated migration rate of core and non-core taxa.

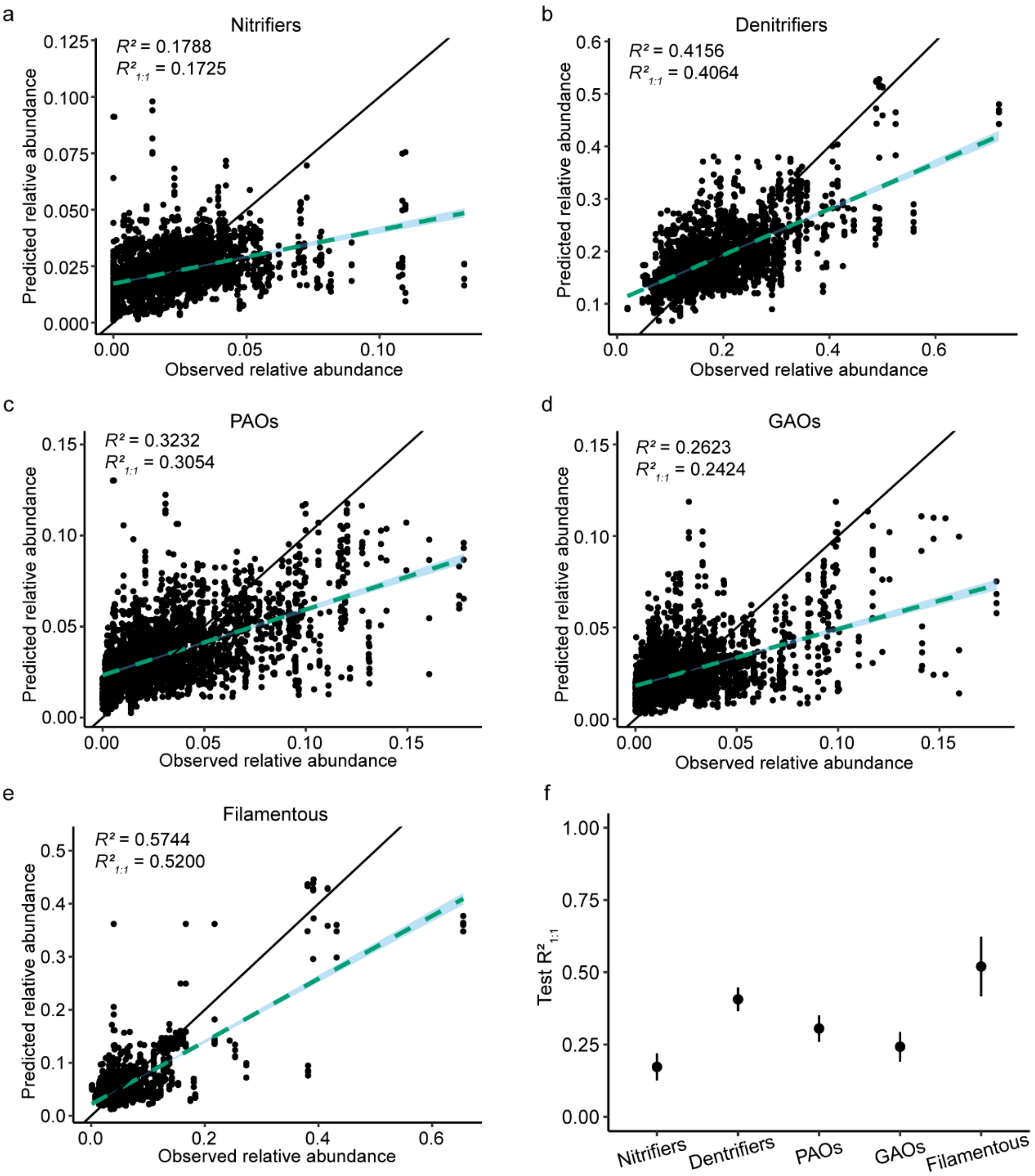


**Fig. S11** Prediction of functional groups with 7 high-weight environmental factors. Correlations between observed and predicted values of nitrifiers (**a**), denitrifiers (**b**), PAOs (**c**), GAOs (**d**), and Filamentous organisms (**e**). **f.** The test $R^2_{1:1}$ of nitrifiers, denitrifiers, PAOs, GAOs, and filamentous organisms.

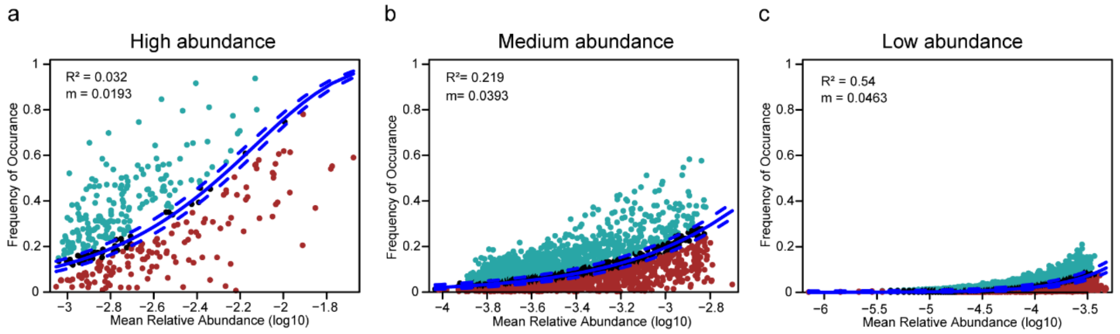


**Fig. S12** Fit of the neutral community model (NCM) of high abundance (**a**), medium abundance (**b**), and low abundance (**c**) subcommunities.

### S3. Supplementary Tables

**Table S1. Abbreviations, meanings and typesof environment variables.**

This table is available as a supplementary dataset; Table S1.xlsx.

**Table S2. Numerical and normalized environmental data.**

This table is available as a supplementary dataset; Table S2.xlsx.

**Table S3. Alpha-diversities of AS system.**

This table is available as a supplementary dataset; Table S3.xlsx.

**Table S4. Summary of ASVs belonging $ASVs_{>10\%}$ subcommunity.**

This table is available as a supplementary dataset; Table S4.xlsx. This table contains the taxonomy, predictive accuracy, occurrence frequency, mean relative abundance, standard deviation, standard error of the mean, coefficient of variation, groups, and partitions of each ASVs in $ASVs_{>10\%}$ subcommunities.

**Table S5. Summary of microbial taxa at different taxonomic levels.**

This table is available as a supplementary dataset; Table S5.xlsx. This table contains the taxonomy, predictive accuracy, mean relative abundance, and occurrence frequency of microbial taxa at different taxonomic levels.

**Table S6. Average importance weights of environmental factors in different ASVs predictive models.**

This table is available as a supplementary dataset; Table S6.xlsx.

**Table S7. Summary of different environment variables.**

This table is available as a supplementary dataset; Table S7.xlsx. This table contains the mean, standard deviation of Garson's connection weights, the mean and coefficient of variation of normalized environment variables, the correlation coefficients of environmental variables with $ASVs_{>10\%}$ subcommunities, and their p-values.

**Table S8. Summary of genera belonging to major functional groups.**

This table is available as a supplementary dataset; Table S8.xlsx.

**Table S9. Summary of prediction results of major functional groups.**

This table is available as a supplementary dataset; Table S9.xlsx. This table contains the predictive accuracies, mean relative abundances, standard deviations, standard errors of the mean, and coefficients of variation of major functional groups.
